# Single *Chlamydomonas reinhardi* (*C. reinhardtii*) cell separation from bacterial cells and auto-fluorescence tracking with a nano-sieve device

**DOI:** 10.1101/2020.02.14.949867

**Authors:** Grant Korensky, Xinye Chen, Mengdi Bao, Abbi Miller, Blanca Lapizco-Encinas, Myeongkee Park, Ke Du

**Affiliations:** Department of Mechanical Engineering, Rochester Institute of Technology, Rochester, New York 14623, United States; Department of Biomedical Engineering, Rochester Institute of Technology, Rochester, New York 14623, United States; Department of Chemistry, Dong-A University, Busan 49315, Republic of Korea; Department of Microsystems Engineering, Rochester Institute of Technology, Rochester, New York 14623, United States; School of Chemistry and Materials Science, Rochester Institute of Technology, Rochester, New York 14623, United States

## Abstract

A planar, transparent, and adaptable nano-sieve device is developed for efficient microalgae/bacteria separation. In our strategy, a sacrificial layer is applied in the dual photolithography patterning to achieve a one-dimensional channel with a very low aspect ratio (1:10,000). Microalgae/bacteria mixture is then introduced into the deformable polydimethylsiloxane (PDMS) nano-channel. The hydrodynamic deformation of the nano-channel is regulated to allow the bacteria cells to pass through while leaving the microalgae cells trapped in the device. At a flow rate of 4 μl/min, ~100% of the microalgae cells are trapped in the device. Additionally, this device is capable of immobilizing single cells in a transparent channel for auto-fluorescence tracking. These microalgae cells demonstrate minimal photo-bleaching over 250 s laser exposure and can be used to monitor hazardous compounds in the sample with a continuous flow fashion. Our method will be valuable to purify microalgae samples containing contaminations and study single cell heterogeneity.

## 1. Introduction

Microalgae have attracted great interest in recent years, with potential applications in renewable energy (Mouwad, 2009; Schneider, 2009), environmental monitoring (Del Carlo et al., 2010), food and feed additives (Brayner et al, 2011), and cosmetics (Barros et al., 2015). For example, microalgal biofuel provides a unique solution to the renewable energy problem because microalgae demonstrate many advantages including high growth rate (Wan et al., 2015), similar co-products to conventional fuels after oil extraction (Spolaore et al., 2006), waste-water remediation capability (Cantrell et al., 2008), no need for arable land (Searchinger et al., 2008), and carbon dioxide fixation capabilities (Chisti et al., 2007). Microalgae can perform charge separation and electron transfer processes with sensitivity to heavy metals, pesticides, explosive compounds and volatile organic compounds measured in a variety of manners (Rouillon et al., 2006; Hernandez et al., 2011; Rea et al., 2011; Bhalla et al., 2011; Brayner et al., 2011). This makes photosynthetic microalgae ubiquitous in researching the monitoring of food, agricultural products, and the aquatic environment for hazardous compounds. Some species of microalgae are auto-fluorescent and the presence of these compounds affects their fluorescent emission spectrum (Marty et al., 1992). These mechanisms are used to create a variety of label-free and amplification-free biosensors (Gosset et al., 2018; Han et al., 2019).

In the case of microalgae biotechnology, label-free cell separation is usually performed to achieve highly purified microalgal populations (Godino et al., 2015). It has been shown that contaminations such as bacteria can cause nutrition competition and affect biomass productivity (Lian et al., 2018). Cell separation methods such as centrifugation, filtration, flocculation, ultrasonic acoustics, and fluorescence activated sorting are typically used to eliminate contaminations (Heaney et al., 1977; Al-Hothaly et al., 2015; Van den Hende et al., 2011; Bosma et al., 2003; Sensen et al., 1993). However, these traditional cell separation methods have several limitations such as cell damage, labor-intensive processing, high expense, difficulty in scaling, and contamination from reagents used in processing. Microfluidic devices utilizing inertia-based separations have been introduced with the advantages such as low costs, small size, rapid response, and high separation efficiency (Dewan et al.,2012; Honsvall et al., 2016). They offer continuous and label-free cell separation, and their ease of use eliminates the need for skilled laboratory technicians to purify microalgal populations through passive filtration. However, these techniques require sophisticated microchannel design and characterization to regulate the flow path of microalgae and bacteria.

We previously introduced a deformable nano-sieve device that can selectively trap microparticles by controlling the hydrodynamic deformation of the shallow channel (Chen et al., 2019; Chen et al., 2020). Furthermore, the captured targets can be easily released by inducing a large deformation of the adaptable channel roof. Leveraging the deformable microfluidics in the present contribution, we show that this device can efficiently and continuously separate microalgae (*Chlamydomonas reinhardii*: *C. reinhardtii*) and bacteria (*Escherichia coli: E. coli*) by controlling the deformed channel height to be larger than the *E. coli* cells but smaller than the *C. reinhardtii* cells without using complicated external actuation. Building upon an integrated fluorescence sensing platform, the technique also enables efficient single cell trapping and analysis by regulating the flow rate and can be used to study single cell behavior and intercellular interactions with controllable cell density. We use this system to monitor the auto-fluorescence signal of *C. reinhardtii* cells over a long period of time. Surprisingly, the auto-fluorescence intensity of *C. reinhardtii* cells only decrease 18% after a 250 s laser exposure, indicating very stable cellular activities. We conclude that our nano-sieve device is a valuable platform for analyzing single microalgae cells with various cellular populations.

## 2. Experiments

### 2.1 Nano-device fabrication

The nano-sieve device was fabricated on a glass substrate. Briefly, 200 nm of tetraethyl orthosilicate (TEOS) was deposited on a 6 inch glass wafer (University Wafer, Inc., MA, USA) by using Plasma Enhanced Chemical Vapor Deposition (AME P5000). We first coated an adhesion promoter (hexamethyldisilazane, HMDS) on the TEOS layer, followed by the spin coating of ~1 μm thick positive photoresist (PR, AZ Mir™ 701) on the substrate. Then, soft bake was applied at 95 °C for 60 s, followed by standard photolithography patterning. After that, the wafer was baked at 100 °C for 60 s and developed by using CD-26 developer (Rohm and Haas Electronic Materials LLC, MA, USA) for 1 min. Buffered Oxide Etching (BOE) was used to etch TEOS layer with an etching rate of ~163.2 nm/min. The sample was immersed into acetone for 1 min to completely remove the PR layer, followed by isopropyl alcohol (IPA) rinsing and cleaning for 15 - 20 s. Finally, the sample was dried by nitrogen gas. To avoid roof collapsing, a second PR layer was patterned on the etched nano-sieve channel by standard photolithography. To seal the channel, Polydimethylsiloxane (PDMS, SYLGARD™ 184, Krayden Inc., CO, USA) base and curing agent were mixed with a ratio of 10:1 and casted to a final thickness of ~3 mm. The PDMS layer was cured at 100 °C for 45 min. The holes on PDMS were punched using a biopsy punch (Miltex Biopsy Punch,1 mm in diameter). Eventually, the PDMS layer was permanently bonded on the glass substrate via oxygen plasma treatment (Electro-Technic Products Inc., IL, USA).

### 2.2 C. reinhardtii cell culture

Bacteria-free *C. reinhardtii* cells and Chlamydomonas culture media were acquired from Carolina Biological Supply (Burlington, NC). The medium inoculated with *C. reinhardtii* cells was vortexed for 5 min to ensure uniform cell distribution. After that, *C. reinhardtii* cells were transferred to a separate vial which was previously sterilized with 100% ethanol and washed thoroughly with Deionized (DI) water. The *C. reinhardtii* cells were stored under a white fluorescent light (~40 W/cm^2^) at room temperature with the cap of the vial loosened to maintain air flow but avoid contamination.

### 2.2 E. coli culture and staining

*Escherichia coli* (*E. coli*, ATCC 25922) cells were cultured at 37 °C for 14-15 hrs, until they achieved an optical density (OD) of 0.5-0.6 (absorption peak: 600 nm). This OD number corresponds to a concentration of ~1.71 × 10^8^ cells/mL. *E. coli* cells were then stained with fluorescent dye for imaging. Briefly, 1 mL of *E. coli* sample was centrifuged at 2000 g for 5 min. The supernatant was then replaced by fresh PBS buffer. We added 4 μL of the BacLight dye into PBS-based *E. coli* solution, followed by vortexing for 10 - 15 s. The stained *E. coli* cells were incubated at room temperature for 20 min and centrifuged again for 5 min, followed by the depletion of the supernatant. Finally, the *E. coli* cells were re-suspended in 500 μL PBS, ready for on-chip experiments.

### 2.3 C. reinhardtii cell auto-fluorescence measurement

Three hundred microliter of *C. reinhardtii* cells were transferred into a 1 mL Eppendorf tube and vortexed for 30 s to further distribute the cells throughout the media. Then, we transferred 100 μL of the cells from the tube to a cuvette and used JASCO FP-8500 Spectrofluorometer to check the fluorescence intensity of the stock solution.

### 2.4 C. reinhardtii cell trapping

We pumped 200 μL of *C. reinhardtii* cells into the nano-sieve device by using a syringe pump (WPI SP220I) with a flow rate ranging from 4 - 8 μL/min. The filtered supernatant was collected in an Eppendorf tube and the auto-fluorescence intensity was measured immediately using Spectrofluorometer. The signal difference between the stock solution and the filtered supernatant was used to calculate the capture efficiency. The nano-sieve channel was also imaged under an Amscope XD-RFL microscope (Irvine, CA) at 4x magnification to determine the overall depositing profile of the *C. reinhardtii* cells. The concentrated area of the deposited *C. reinhardtii* cells was imaged at 40x magnification.

### 2.5 Auto-fluorescence tracking

The auto-fluorescence signal of the *C. reinhardtii* cells trapped in the nano-sieve device was tracked by using a sensitive benchtop fluorometer (Qin et al., 2019), excited with a continuous wavelength laser (488 nm, 2 mW). The signal was collected by using a parabolic mirror, filtered by a notch filter, and detected by using a portable spectrometer (USB 2000+).

### 2.6 SEM imaging

The fabricated nano-sieve channel was imaged by using scanning electron microscopy (SEM, Tescan Mira3). A thin metal film (~5 nm) was coated on the surface of the sample by using SPI-Module™ Sputter Coater. For high resolution imaging, the applied voltage was set at 20 kV.

## 3. Results

The schematic of the experimental design is shown in **Fig. 1a**: *C. reinhardtii* cells were pumped into a shallow and wide nano-sieve channel with a height of ~200 nm. Since the diameter of the *C. reinhardtii* cells is ~10 μm, they were trapped in the shallow channel as a monolayer, allowing cell media to be pumped to the outlet. The micrograph of the nano-sieve device is shown in **Fig. 1b**. Each channel has a dimension of 2 mm by 12 mm, which enables to pattern ~30 channels on a single wafer for high throughput experiments. Before experiments, channels were filled with blue food dye to confirm there were no collapsing issues. As shown in **Figs. 1c-i**, before on-chip experiments, original *C. reinhardtii* cells show light green color (white dashed box). On the other hand, the collected supernatant solution from the nano-sieve outlet shows transparent color, even though the volume does not have significant change. We used microscope to image the original solution and filtered supernatant, which are presented in **Fig. 1c-ii** and **1c-iii**, respectively. Compared with original solution, the filtered solution only has very few cells, demonstrating that our device can efficiently trap *C. reinhardtii* cells. The *C. reinhardtii* cells trapped within the nano-sieve device were also imaged by the microscope. As shown in **Fig. 1d**, the *C. reinhardtii* cells are uniformly patterned in a parabolic shape (dashed white line), which is in agreement with our deformation theoretical model (Chen et al., 2019).

**Figure 1.**
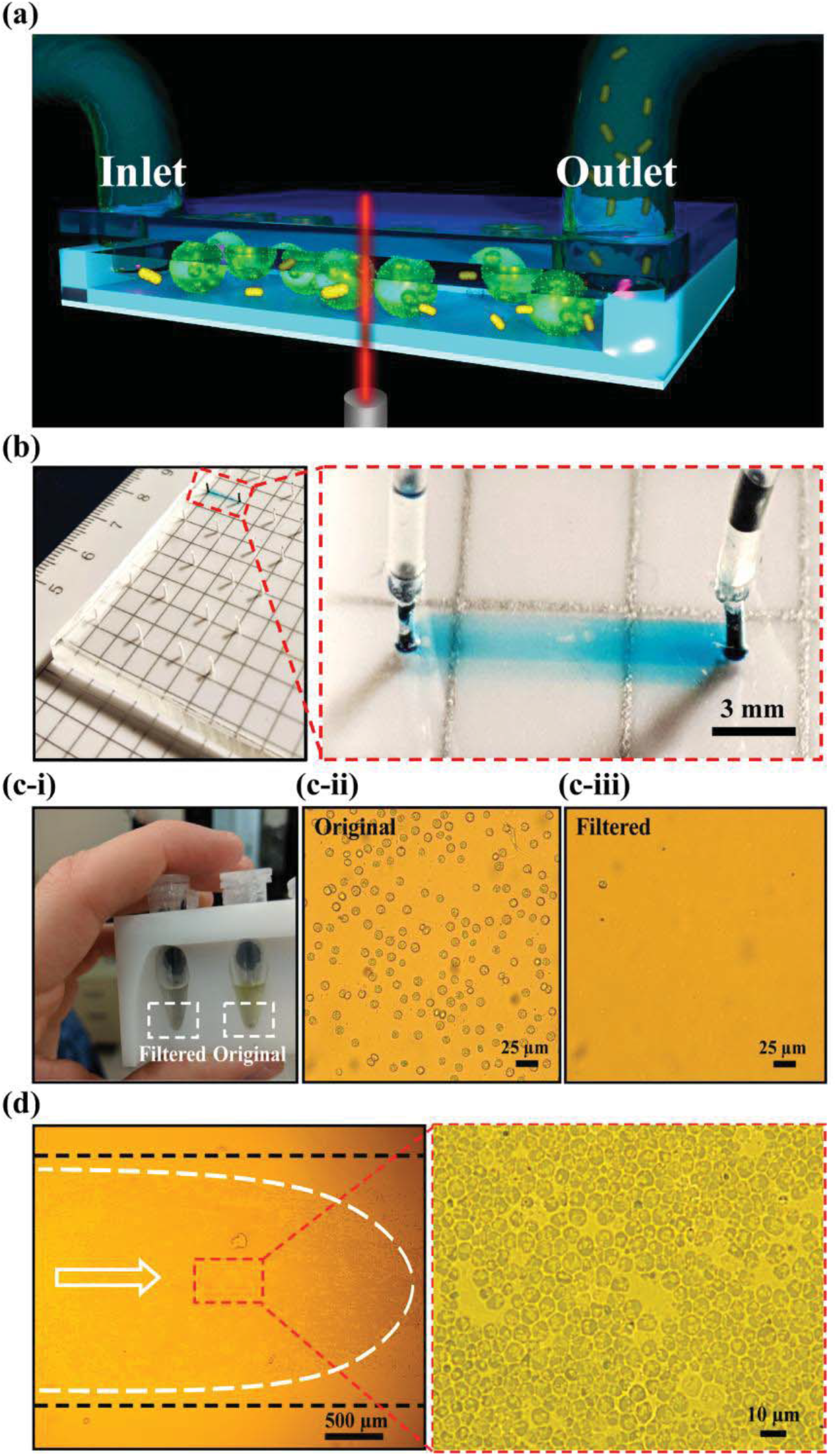
(a) Schematic of experimental design for nano-sieve device trapping *C. reinhardtii* cells; (b) The optical micrograph exhibiting the practical device (left); the blue food dye is filled within the entire channel (right); (c-i) The filtered solution and original solution collected in the centrifuge tube. The microscope images showing the density of cells before (c-ii) and after (c-iii) the filtering process. (d) The *C. reinhardtii* cells trapped within the nano-sieve channel. The white arrow indicates the flow direction. The white dashed parabola shows the profile of the trapped cells. The enlarged region shows the dense *C. reinhardtii* cells.

We developed a double photolithography patterning protocol to avoid channel collapsing during channel sealing. The fabrication procedure is shown in **Fig. 2a**. First, the channel was patterned via standard photolithography and TEOS wet etching on a glass substrate. After dissolving the PR layer, we patterned a second PR layer in the channel area only. Thus, after plasma treatment on the PDMS and TEOS layer, the channel was sealed with permanent bonding except at the area coated with the second PR layer. After channel sealing, the second PR layer was easily removed by filling acetone in the channel. The channels patterned with and without the second PR layer are shown in **Fig. 2b-i** and **2b-ii**, respectively. A rectangular red shape indicates the presence of the PR layer. The SEM image of the channel patterned with PR is shown in **Fig. 2c**, confirming the PR layer was patterned in the center of the nano-sieve channel. If the second PR layer is not utilized, then the channel is easily blocked with the PDMS collapsing on the substrate. We used an integrated pressure control system (Precigenome LLC, CA, USA) to monitor the air pressure in the nano-sieve. As shown in **Fig. 2d**, the air flow rate in the nano-sieve device increases significantly (~20 to ~100 sccm) in the absence of PR (blue). In comparison, the measured air flow rate for channels patterned with PR only reaches to ~2.3 sccm (red). Thus, the air pressure measured in the channel without the second PR layer was ~50 times higher than the channel patterned with the second PR layer, indicating a very high flow resistance due to channel collapse.

**Figure 2.**
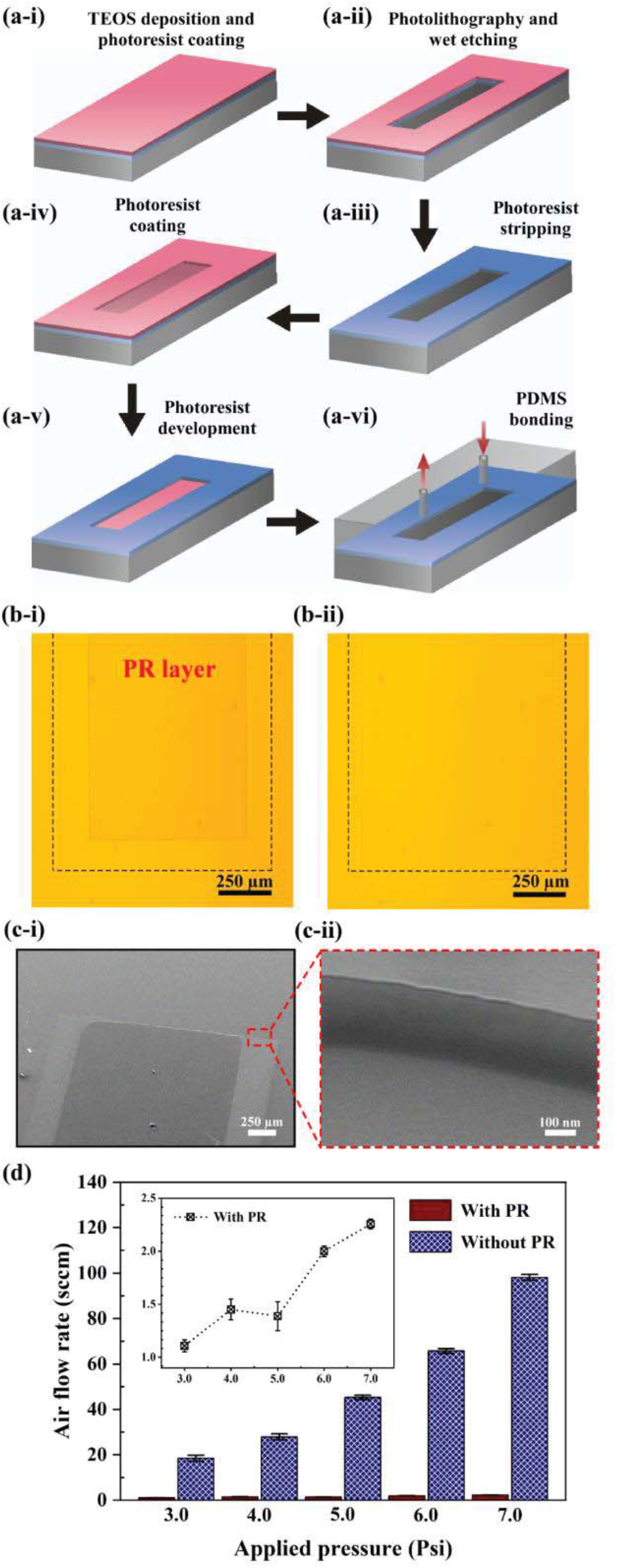
(a) The fabrication process of nano-sieve device by using dual photolithography. (b) Microscope images of the sacraficial PR layer: (b-i) before and (b-ii) after acetone washing. (c) SEM image of (c-i) low maginication and (c-ii) high magnification of the scarifical PR layer. (d) The measured air flow rate vs. applied pressure in the nano-sieve with (red bar) and without (blue patterned bar) PR layer.

The *C. reinhardtii* cell capture efficiency vs. applied flow rate is shown in **Fig. 3a**. The auto-fluorescence intensities of original and supernatant solution were compared by using a spectrofluorometer with an excitation wavelength of 488 nm. With a flow rate of 4 μL/min, most of the *C. reinhardtii* cells were trapped in the nano-sieve device as the fluorescence intensity dropped from ~13 counts to ~3 counts (**Fig. 3a-i**). Increasing the flow rate decreases the cell capture efficiency. For example, with a flow rate of 6 μL/min, the fluorescence intensity dropped from ~13 counts to ~5 counts, indicating some of cell leaking (**Fig. 3a-ii**). At a flow rate of 8 μL/min, the supernatant intensity reached to 15 counts and is comparable with the stock solution (**Fig. 3a-iii**), demonstrating most of the cells were washed to the outlet. Our device can also be used to separate microalgae and bacteria sample. We pumped labeled *E. coli* and *C. reinhardtii* cell mixture into the nano-sieve device. As shown in **Fig. 3b**, the mixture solution shows two distinct fluorescence peaks: the labeled *E. coli* cells emit at 520 nm and the *C. reinhardtii* cells emit at 680 nm. After on-chip experiments, almost all the *E. coli* cells passed through the nano-sieve with a flow rate ranging from 4 to 8 μL/min (**Fig. 3b-i to Fig. 3b-iii**). However, the *C. reinhardtii* cells were trapped in the device as the supernatant without showing significant fluorescence peak at 680 nm.

**Figure 3.**
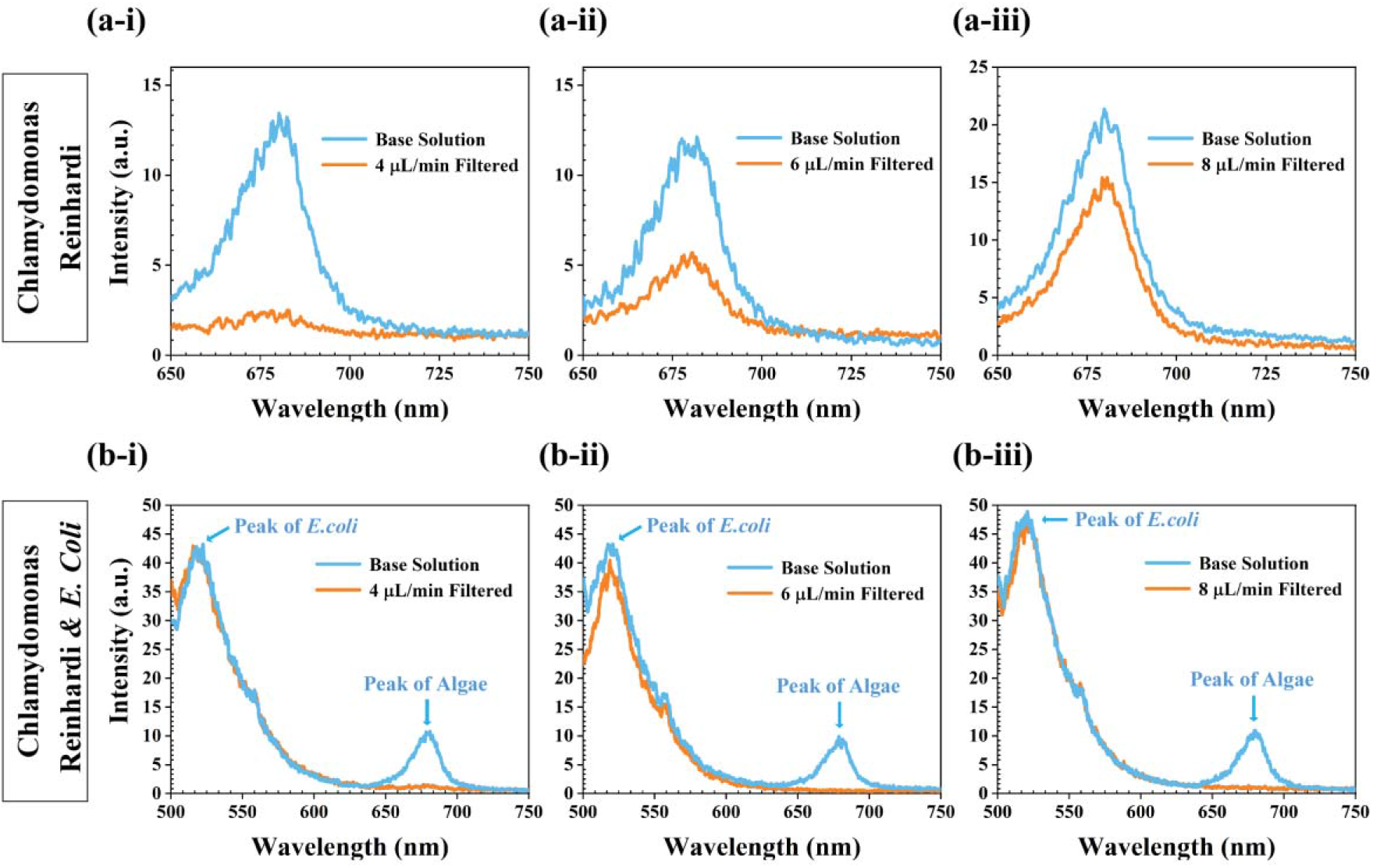
(a) The *C. reinhardtii* cells capture efficiency vs. various applied flow rates of (a-i) 4 μL/min; (a-ii) 6 μL/min; (a-iii) 8 μL/min, respectively. (b) The mixture of *E. coli* and *C. reinhardtii* cell separated by nano-sieve device, under the applied flow rate from (b-i) 4 μL/min; (b-ii) 6 μL/min; (b-iii) 8 μL/min, respectively. The blue curve is the original solution and the orange curve is the filtered supernatant.

The nano-sieve device can be used to track the auto-fluorescence emission of microalgae under mechanical and chemical stresses. To show this, we mounted the nano-sieve device onto a benchtop fluorometer with a laser wavelength of 488 nm (2 mW). The laser beam width is ~800 μm, thus we can excite ~6000 cells collectively. The auto-fluorescence signal from the *C. reinhardtii* cells was collected by using a parabolic mirror, installed under the nano-sieve chip. An optical fiber was used to collect the signal and the signal was analyzed by a small spectrometer (**Fig. 4a**). We measured the auto-fluorescence intensity every 50 s with a total time of 250 s. As shown in **Fig. 4b**, the auto-fluorescence intensity of the *C. reinhardtii* cells was very stable as it only changed from 7.7 ×10^4^ to 7.6 ×10^4^ counts from 0-50 s; 7.6×10^4^ to 7.3×10^4^ counts from 50-100 s; 7.3 ×10^4^ to 7.0 ×10^4^ counts from 100-150 s; 7.0 ×10^4^ to 6.7 ×10^4^ counts from 150-200 s; and 6.7 ×10^4^ to 6.3 ×10^4^ counts from 200-250 s. Over the duration of 250 s laser excitation, the auto-fluorescence intensity only dropped 18%, indicating this set-up can be used for single cell monitoring.

**Figure 4.**
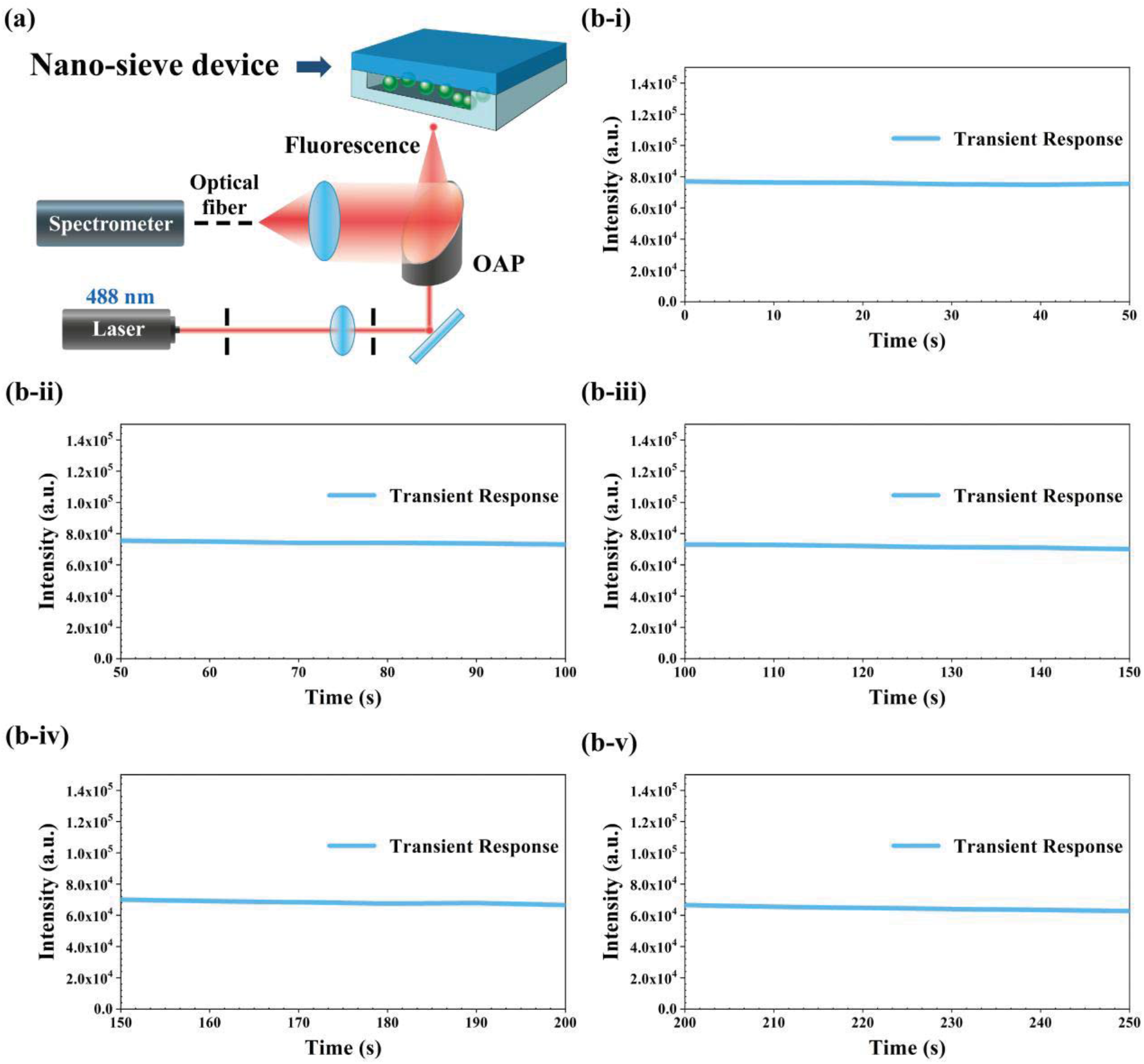
(a) The integrated fluorometer used to measure the auto-fluorescence signal from the trapped *C. reinhardtii* cells. (b) The auto-fluorescence intensity of *C. reinhardtii* cells under continuous excitation: (b-i) 0-50 s; (b-ii) 50-100 s; (b-iii) 100-150 s; (b-iv) 150-200 s; (b-v) 200-250 s.

## 4. Discussion

Our nano-sieve device is capable of immobilizing single cells in a transparent and planar environment and can be used to study their heterogeneous phenotypes (Thibault et al., 2019; Prakadan et al., 2017). Other approaches, such as T junctions (Jin et al., 2015) and microstencils (Zaidi et al., 2018) have been added into microfluidic devices to immobilize single cells. However, all these approaches rely on complicated micro/nanofabrication methods and are sensitive to the cell density in the stock solution. Our cell trapping methodology relies on a wide and shallow channel and is achieve by continuous flowing the cells into the channel without clogging/loss issues. As the channel height and deformation of the nano-sieve can be tuned, we should be able to separate and study different cell types with various sizes. As shown in **Fig. 1**, the trapped *C. reinhardtii* cells has a larger fringe than the cells presented in cell media, indicating that the cells were deformed by the flexible PDMS roof. Thus, our device also has potential for quantitative studies of cell mechanics by controlling the PDMS deformation (Mills et al., 2008; Huh et al., 2007; Du et al., 2017). This configuration can also be applied to study the metabolic and physiological responses of the trapped microalgae cells to the external chemical/mechanical triggering (Ferraria et al., 2019; Liu et al., 2013).

One of the main challenges of the fabrication of shallow nano-sieve device is the bonding between the PDMS roof and the glass substrate. We used a positive PR layer as a sacrificial layer to cover the etched nano-sieve channel. Thus, oxygen plasma only generated the hydroxyl functional group on the substrate outside of the etched channel. After baking to seal the channel, solvent such as acetone was used to easily dissolve the sacrificial layer. By using this simple approach, we are able to pattern nano-sieves with an extremely low aspect ratio of 1:10,000 (height: width) without roof collapsing issues. As shown in **Fig. 2d**, without roof collapsing, the measured air flow pressure is as low as 2.3 sccm, indicating very low resistance built in the channel. This type of extremely low aspect ratio nano-device can be further used for microparticle and pathogen separation (Chen et al., 2020), droplet microfluidics (Ying et al., 2013), and reconfigurable optofluidics (Brennan et al., 2009).

To explore microalgae as the next generation biofuel, it is important to keep microalgae population viable over a long period (Brayner et al., 2011). Microalgae populations can be adversely affected by bacteria populations and other forms of biological contaminants. For example, certain microbes can damage the microalgae cell wall and prevents its growth (Afi et al., 1996; Arora et al., 2012; Wang et al., 2015). We show that our deformable nano-sieve device can efficiently separate microalgae from bacteria, enabling controlled movement or trapping of microalgae while mitigating the presence of contaminants. Even though other microfluidic methods have been introduced for microalgae and bacteria separation, they typically require complicated microchannel design and characterizations of the fluid elasticity (Dewan et al., 2012; Hønsvall et al., 2016; Jeonghun et al., 2019). In our case, the separation is achieved by fine-tuning the flow rate and hydrodynamic deformation of the nano-sieve. As shown in **Fig. 3b**, almost all the contaminant bacteria cells were washed to the waste reservoir without showing loss of microalgae. However, micrographs taken before and after the mixed *C. reinhardtii* and *E. coli* solutions indicate that the amount of *C. reinhardtii* cells trapped in the channel was lower than expected, even though a significant amount of *C. reinhardtii cells* was still visible in the channel. This indicates that a portion of the *C. reinhardtii* cells was lysed when pumped through the channel with *E. coli* cells, proving that the *C. reinhardtii* cell membrane may be destroyed by the presence of *E. coli*.

In this work, we have taken the first step to establish a reliable, inexpensive, and label-free separation and biosensor method. Furthermore, stable sensing can be implemented since, as shown in **Fig. 4**, *C. reinhardtii* cells can withstand continuous laser excitation for minutes with minimal photobleaching issues. Many of the microalgae biosensors constructed to date are limited by the stability of the fluorescence signal to be measured over time and various immobilization techniques are being investigated to solve this problem (Brayner et al., 2011). Large fluctuations in the auto-fluorescent signal of the *C. reinhardtii* cells could lead to false-positive and false-negative results. Combining the small and sensitive fluorometer with *C. reinhardtii* cells trapped nano-sieve device, we demonstrated excellent auto-fluorescence stability of *C. reinhardtii* cells over a long period, which can be further used for rapid and label-free detection of aquatic pollutants (Gosset et al., 2018). The proposed method can also be used for single-cell analysis of genomics, transcriptomics, and metabolomics without involving complicated serial dilution, micromanipulation, and dielectrophoretic sorting.

## 5. Conclusions

In summary, we have demonstrated a novel nano-sieve device for efficient and simple microalgae/bacteria separation at the single-cell level using controllable hydrodynamic deformation. The utilization of a sacrificial layer avoided channel collapsing and leaking issues while minimizing the flow resistance for the separation process. Taking advantage of deformable microfluidics, single *C. reinhardtii* cells were trapped in the nano-device and subsequently separated from *E. coli* cells. This nano-sieve device was also integrated with a small fluorescence sensing unit to continuously track the change in auto-fluorescence over time. With minimal photo-bleaching issues, our method has the potential to be used as a label-free biosensor for environment monitoring. Future work will extend this method for multiplexing and high throughput bio-and chemical sensing.

## Declaration of interests

⊠The authors declare that they have no known competing financial interests or personal relationships that could have appeared to influence the work reported in this paper.

□The authors declare the following financial interests/personal relationships which may be considered as potential competing interests:

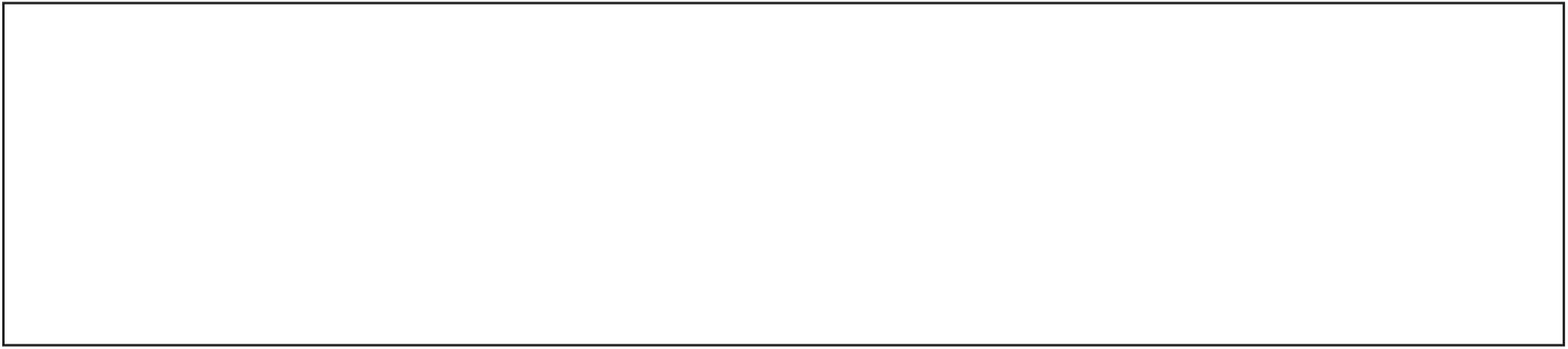

